# SARS-CoV-2 occurrence in white-tailed deer throughout their range in the conterminous United States

**DOI:** 10.1101/2023.04.14.533542

**Authors:** Sarah N. Bevins, Richard B. Chipman, Scott F. Beckerman, David L. Bergman, Derek T. Collins, Thomas J. Deliberto, Joshua P. Eckery, Jeremy W. Ellis, Allen L. Gosser, Jonathon D. Heale, Jason M. Klemm, Kristina Lantz, Timothy J. Linder, Mitch Oswald, Robert Pleszewski, Christopher A. Quintanal, Jourdan M. Ringenberg, Kelsey R. Weir, Mia K. Torchetti, Julianna B. Lenoch, Jeffrey C. Chandler, Susan A. Shriner

## Abstract

Broad-scale data show SARS-CoV-2 occurrence in white-tailed deer throughout much of their range in the conterminous United States and reinforce findings of considerable SARS-CoV-2 infection and exposure. Results shed light on both current infections and prior exposure, with prevalence decreasing over time and seroprevalence increasing.

**One-Sentence Summary:** White-tailed deer are infected with, and have been exposed to, SARS-CoV-2 throughout their range in the conterminous US.

## Main Text

There is now clear evidence that SARS-CoV-2 can move from humans to multiple other animal species (1). This is especially true for white-tailed deer (*Odocoileus virginianus*) and possibly other cervid species in North America, where early research found 40% of white-tailed deer had been exposed to SARS-CoV-2 (2) and more than one-third of deer in some populations were positive by rRT-PCR (real-time PCR with reverse transcription) (3). This is an unexpectedly high level of infection for a recently emerged pathogen than has now spilled-over from people to wildlife. Comparison of deer and human lineages suggests human-to-deer transmission is occurring repeatedly, followed by deer-to-deer transmission (4) which creates the potential for cervids to become a reservoir for a pathogen that can then spillback into the human population. Widespread persistence of the virus in multiple species exposes the pathogen to varying selection pressures, potentially leading to evolution of novel variants (5, 6).

Given these concerns, a national-scale monitoring effort was initiated to monitor both viral infection and antibodies indicative of SARS-CoV-2 exposure in populations of white-tailed deer across the United States (US) (7, 8). Broad-scale surveillance helps illuminate differences in infection and exposure across geographic regions and habitat types. In addition, capturing both prevalence and seroprevalence in combination helps to clarify the extent of current and prior exposure, providing insight into viral dynamics.

## Methods

Oral or nasal swabs preserved in PrimeStore Molecular Transport Media (MTM, Longhorn Vaccines and Diagnostics LLC), along with whole blood samples collected on Nobuto filter paper, were collected from white-tailed deer from 29 states throughout their range in the US (Fig 1). The majority of samples were collected in collaboration with tribal and state wildlife management agencies from hunter-harvested white-tailed deer, with additional samples coming from roadkill and deer being removed as part of permitted wildlife damage management activities. Sampling was concentrated in fall and winter when hunter harvest activities typically occur, although roadkill and wildlife damage management samples were collected year-round. Blood samples on Nobuto filter paper from Ohio were provided by the Ohio Animal SARS-CoV-2 Surveillance Consortium (Ohio State University Department of Veterinary Preventive Medicine, Ohio Department of Natural Resources Division of Wildlife, and US Department of Agriculture Wildlife Services).

**Figure 1.**
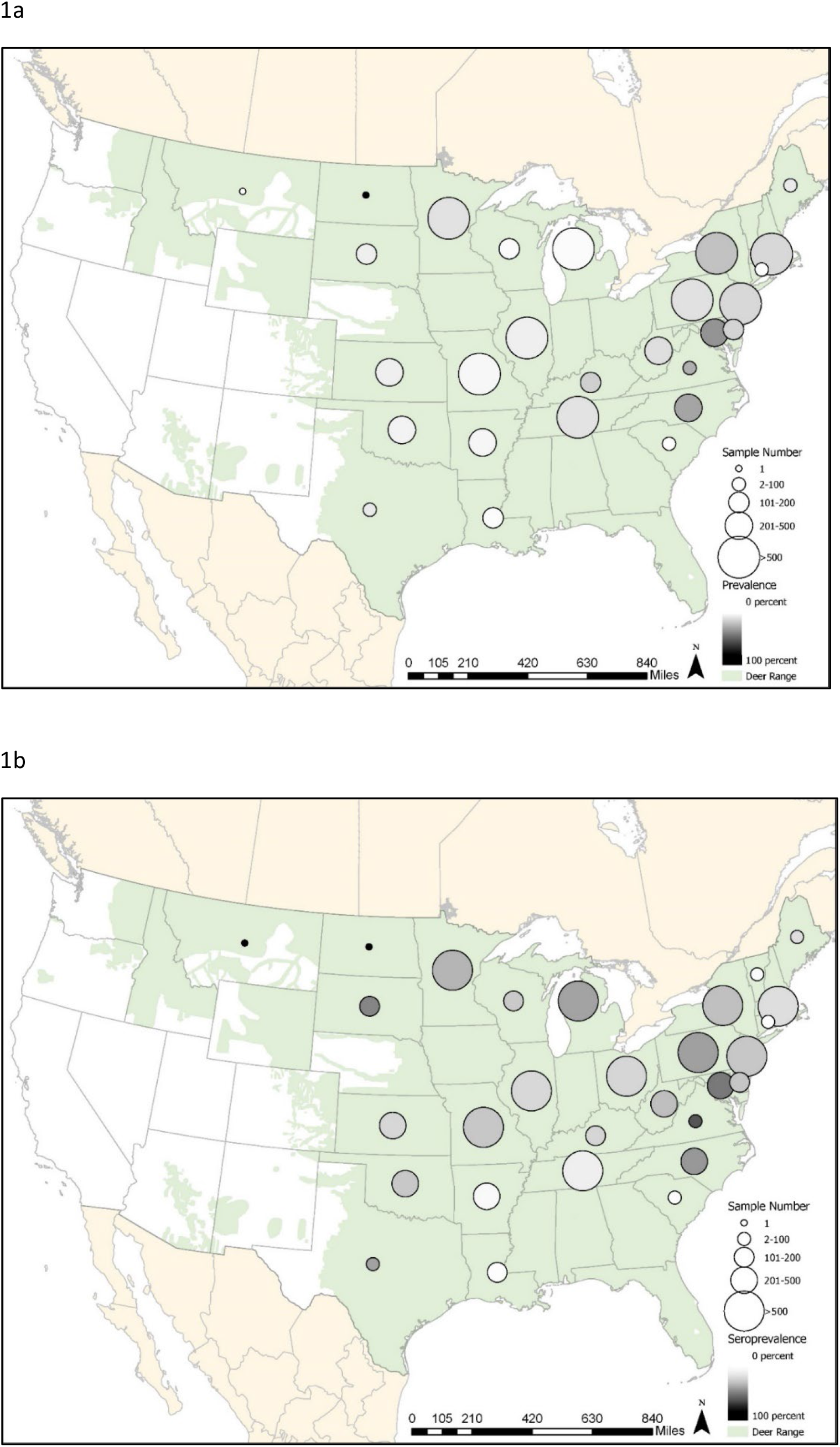
a) Infection prevalence and b) seroprevalence of SARS-CoV-2 in white-tailed deer. Circle size indicates the relative number of samples tested and gray shading indicates the relative seroprevalence and rRT-PCR prevalence in a state. White-tailed deer distribution in the conterminous US is shown in green. White-tailed deer range data from Hanberry, Brice B.; Hanberry, Phillip. 2019. Digitized white-tailed deer densities for the continental United States. Fort Collins, CO: Forest Service Research Data Archive.https://doi.org/10.2737/RDS-2019-0053

SARS-CoV-2 RNA was extracted from oral and nasal swab samples using MagMAX™ CORE Nucleic Acid Purification Kits (Applied Biosystems) in accordance with the manufacturer’s instructions. Singleplex rRT-PCR of SARS-CoV-2 N1 and N2 was performed using 5 μL of extracted RNA with the BioRad Reliance

One-Step Supermix kit. SARS-CoV-2 Research Use Only qPCR Primers & Probes for N1 and N2 were provided by Integrated DNA Technologies. Thermal cycling conditions were 50°C for 10 min and 95°C for 10 min followed by 45 cycles of 95°C for 10 sec and 60°C for 30 sec using a BioRad CFX96 Touch Real-Time PCR Detection System or CFX Opus Real-Time PCR System. rRT-PCR results from swab samples collected in Ohio was previously reported (3).

Antibodies were extracted from Nobuto filter paper strips as described previously (9) and screened at a functional dilution of 1:20. Extracted samples were screened using a surrogate virus neutralization test (sVNT, Genscript cPass™) using a VarioSkan Flash or Varioskan LUX multimode microplate reader (Thermo Fisher). At least two technical replicates were used to calculate an average % inhibition. The sVNT has not been validated for deer; however, previous evaluations with white-tailed deer sera samples suggested that sVNT results were qualitatively similar to a highly specific SARS-CoV-2 virus neutralization test (VNT)(2).

## Results

During November 2021 to October 2022, we collected 11,113 white-tailed deer samples from 29 states and Washington, DC. Samples with incomplete data were omitted from analysis, and overall, SARS-CoV-2 was detected in 12.2% (1,116 out of 9,091 tested) of white-tailed deer sampled and 31.6% (3,363 out of 10,639 tested) had antibodies indicative of SARS-CoV-2 exposure.

While well-sampled (more than 300 samples) states exhibited a wide range in prevalence (1.3% in Michigan to 35.2% in North Carolina) (Fig 1a), serological exposure was typically high (Fig. 1b), and as expected, higher than viral prevalence. The lowest seroprevalence rate in states with more than 300 samples was found in Tennessee, where 13.1% of white-tailed deer had evidence of SARS-CoV-2 exposure (mean = 32.1% in states with more than 300 samples) (Fig. 1b). Sample collection primarily occurred between November 2021 through April 2022; prevalence declined over time, while seroprevalence gradually increased (Table 1).

**Table 1:** Prevalence, seroprevalence, and sample sizes over time.

| Month | rRT-PCR<br>Prevalence | # rRT-PCR<br>Tested | sVNT<br>Seroprevalence | # sVNT<br>Tested |
| --- | --- | --- | --- | --- |
| November-21 | 19.86 | 1465 | 28.72 | 2347 |
| December-21 | 16.21 | 2381 | 25.44 | 2492 |
| January-22 | 9.23 | 1582 | 36.5 | 1907 |
| February-22 | 10.78 | 1883 | 35.88 | 2051 |
| March-22 | 5.76 | 1232 | 35.36 | 1346 |
| April-22 | 10.46 | 153 | 43.52 | 108 |
| May-22 | 0 | 10 | 50 | 4 |
| June-22 | 0 | 1 | -- | -- |
| July-22 | 0 | 32 | 32.35 | 34 |
| August-22 | 0 | 127 | 42.15 | 121 |
| September-22 | 0 | 149 | 23.33 | 150 |
| October-22 | 0 | 76 | 1.27 | 79 |

Analysis of white-tailed deer that were simultaneously assayed for both infection and serology revealed that 23.3% (1,995/8,542) were rRT-PCR negative but antibody positive, an indication of widespread prior infection (2). In addition, 4.1% (354/8,542) were positive by rRT-PCR but not serology, likely indicating recent or first-time infections. In instances where rRT-PCR positive samples also had a corresponding blood sample, more than half (54.8%, 519/947) were also serologically positive.

## Discussion

These broad-scale data show SARS-CoV-2 occurrence in white-tailed deer throughout the majority of their conterminous US range and reinforce findings of considerable SARS-CoV-2 infection and exposure identified in prior, more localized studies. (2–4).

While infection was widespread, findings also show a decrease in prevalence over the course of the study. The overall prevalence reported here (12.2%) is also lower than smaller-scale white-tailed deer studies completed recently (Ohio=35.8%, Iowa=33.2%). These differences may simply reflect spatial heterogeneities of disease spread at small versus large scale or could be related to a variety of factors such as seasonal epidemiological dynamics that occurred during the course of the individual studies. It is also possible that the dominant Omicron variant currently circulating in human populations is less likely to infect white-tailed deer compared to earlier SARS-CoV-2 variants. Indeed, experimental work showed that the Omicron variant (B.1.1.529) was less likely to infect transgenic mice and caused less disease in wild-type hamsters compared to previous variants (8, 10). If this is the case, viral dynamics in white-tailed deer could shift from recurring spillover from human populations and instead be driven by deer-to-deer transmission. Regardless, the differences likely demonstrate what will continue to be a shifting landscape in the dynamics of SARS-CoV-2 in white-tailed deer and highlight the importance of continued long-term monitoring.

The high seroprevalence (31.6%) findings in comparison to rRT-PCR positives (12.2%) is not a surprise given the longer time horizon for antibody detection, but how long the antibodies persist is not clear. Long-term studies of SARS-CoV-2 in white-tailed deer have not been carried out, although some studies suggest antibodies can persist up to 13 months in captive white-tailed deer (11, 12). For samples where both virus and antibodies were detected (5.7%, 519/9,091), findings could be indicative of prior exposure or could suggest that seroconversion happened rapidly, overlapping with the period that virus was still detectable in swabs. Both dynamics–prior infection and rapid seroconversion–are likely occurring. Palmer et al. (2022) showed seroconversion often occurs within one week of infection, with virus still being detectable in some cases up to 21 days later (13).

Simultaneous collection of both rRT-PCR and serology data was the suggested next step from Chandler et al. 2022 for national surveillance, and the implementation and resultant findings reported here provide a full picture of infection and exposure in white-tailed deer, although regional level differences in infections and exposure indicate that the window of infection may have been missed in some cases. Mule deer (*Odocoileus hemionus*) in the US are the only other cervid species confirmed to be infected with SARS-CoV-2 (14), but to date, those findings have been rare. There is also currently no evidence of infections in additional cervid species in other countries where research has been carried out (15), making white-tailed deer unique in the currently available body of research. Broadscale and comprehensive findings like those reported here can also inform downstream research working to identify the transmission pathways moving virus from humans to white-tailed deer and back again (5). Understanding these drivers is the next step in identifying management actions that could lessen spillover from people to wildlife.

## Acknowledgements

We thank Wildlife Services employees and our collaborators at tribal nations and state wildlife agencies for contributing wildlife sampling expertise, as well as hunters for participating in this large-scale effort.

